# Electrode pooling preserves movement decoding by retaining neural population dynamics

**DOI:** 10.64898/2026.05.13.724949

**Authors:** Shih-Hung Yang, Yu-Chun Lin, Wen-Yen Hsieh, Yi-Fang Chen, Wan-Jung Chung, Yu-Shun Liu, Yi-Ko Chen, Yao-Ting Chiu, Shi-Sheng Shen, Yu-Wei Wu

## Abstract

New implantable-electrode fabrication strategies make dense, ultrafine electrode arrays with lower tissue burden increasingly feasible, shifting a key bottleneck for scalable brain-computer interfaces from electrode placement to readout capacity. Electrode pooling, in which multiple electrodes share a readout channel, could relax this bottleneck by combining extracellular signals before acquisition, but it has remained unclear whether such compression preserves the neural population structure needed for behavioral decoding. Here we evaluate this question using software-emulated electrode pooling in mouse sensorimotor cortex during a cue-guided reach-and-grasp task using a high-density microwire array coupled to a CMOS microelectrode-array platform. Pooled recordings retain forelimb kinematic information more effectively than a channel-matched control that discards electrodes. Pooling reduces the separability of electrode-specific spikes and sorted units, indicating a loss of some neuronal detail, but the mixed signals still preserve task-aligned low-dimensional latent dynamics that support decoding. When readout capacity is fixed, this trade-off allows broader electrode coverage to contribute to behaviorally informative population sampling. Together, these results define electrode pooling as a design trade-off for scalable readout, in which some electrode-specific neuronal information is lost but the population dynamics needed for movement decoding remain accessible.

## Introduction

Recording from large neural populations is central to circuit neuroscience and brain-computer interface (BCI) development^1–6^, and advances in high-density silicon probes, CMOS-integrated microwire systems, and large channel-count architectures have greatly expanded the number of implantable recording sites^2–4,7^. Miniature microwires are especially attractive because they support chronic recordings and, at sufficiently small diameters, can be inserted with minimal acute tissue disruption^8,9^. Recent *in vivo* mechanics studies showed that small-diameter microwires can displace rather than rupture cortical blood vessels, while ultrasmall bioactive microelectrodes and flexible arrays further illustrate the growing feasibility of dense microscopic neural interfaces^7–12^. Yet scalable BCI devices remain constrained by the readout electronics that must amplify, filter, digitize and transmit neural signals. In practice, channel budget, power consumption, heat dissipation, interconnect routing and package size often limit how many electrodes can be sampled simultaneously, even when many more electrodes can be fabricated or implanted^5,6^.

Several strategies have been proposed to improve the efficiency of neural interfaces. Time-division multiplexing, electronic depth control and programmable electrode switching allow many physical sites to share fewer readout paths, while CMOS microelectrode arrays integrate dense electrode grids with local electronics to reduce routing overhead^13–19^. Complementary signal-processing and decoding approaches reduce the burden after acquisition by transmitting threshold crossings or features rather than raw voltage traces, by using multiunit or spike-band signals when single-unit isolation is unstable, or by aligning low-dimensional neural activity to maintain decoder performance across sessions^20–23^. These solutions improve different parts of the BCI pipeline, but many still preserve a one-electrode-to-one-channel measurement at any instant or compress signals only after they have already occupied a readout channel.

Electrode pooling provides a more direct form of front-end compression^24^. Lee et al. proposed using controllable switches in silicon probes so that several recording sites can be connected to one wire at the same time, causing their extracellular voltages to be combined before leaving the probe. This architecture differs from static switching, which selects only one site, and from time-division multiplexing, which samples different sites sequentially. Their study implemented pooling on Neuropixels 1.0, characterized the resulting signal and noise, and used simulations to show that, under suitable configurations, pooling can save wires without necessarily compromising the content of extracellular recordings^24^. The central intuition is that extracellular spikes are sparse in time and spatially local, so mixed electrode signals can remain sortable or informative when the pooled electrodes are chosen appropriately^20,25,26^.

However, an important question remained unresolved. The prior pooling framework addressed electrophysiological yield and signal fidelity, but it did not test whether pooled recordings preserve the neural population information needed for behavioral decoding in a motor BCI-like setting. Pooling could be detrimental if voltage mixing removes the temporal structure required to infer movement. Conversely, if task-relevant activity is represented in low-dimensional latent dynamics that are stable to changes in the identities of individual sorted units, then pooling may preserve the most important information even while reducing per-electrode separability^21–23,27–30^. This unknown directly motivates the present study.

Here we tested electrode pooling in mouse motor cortex during a cue-driven reach-to-grasp task using a high-density microwire array interfaced to a CMOS microelectrode array system. This microwire platform allowed us to test whether the biological scalability of low-trauma miniature electrodes can be converted into decoding performance when readout electronics remain fixed. We compared split mode (one electrode per channel), electrode pooling (2-6 electrodes per channel), and a channel-matched random picking control that kept only one electrode per pool. We show that pooling retained strong kinematic decoding with up to 6-fold channel compression, outperformed the channel-matched control, preserved low-dimensional latent dynamics aligned to split mode, and improved decoding when more electrodes were attached to each channel under a fixed channel budget. These results position electrode pooling as a practical strategy for scaling neural interfaces by implanting more low-trauma microwires than can be individually read out while prioritizing task-relevant population dynamics over strict one-electrode-per-channel separability.

## Results

### Electrode pooling retains decoding under 2- to 6-fold channel compression

We first defined the acquisition schemes and behavioral benchmark used throughout the study (**Fig. 1**). In split mode, each electrode was connected to an independent amplifier, filter and ADC chain (**Fig. 1a**). In electrode pooling, multiple electrodes were tied to one readout channel through a pooling adaptor (**Fig. 1b**). As a channel-matched control, we used random picking, in which each predefined pool was reduced to one randomly selected electrode while preserving the same number of output channels and the same spatial tiling as pooling (**Fig. 1c**). This control separates the effect of signal mixing from the effect of channel reduction alone: if decoding were maintained simply by keeping a sparse subset of electrodes, random picking should perform similarly to pooling. We implemented software pooling by combining split-mode traces to emulate the hardware configuration on the CMOS MEA platform (**Fig. 1d**). The implanted mouse performed a cue-guided reach-to-grasp behavior while forelimb kinematics were tracked with high-speed video (**Fig. 1e,f**).

**Fig. 1.**
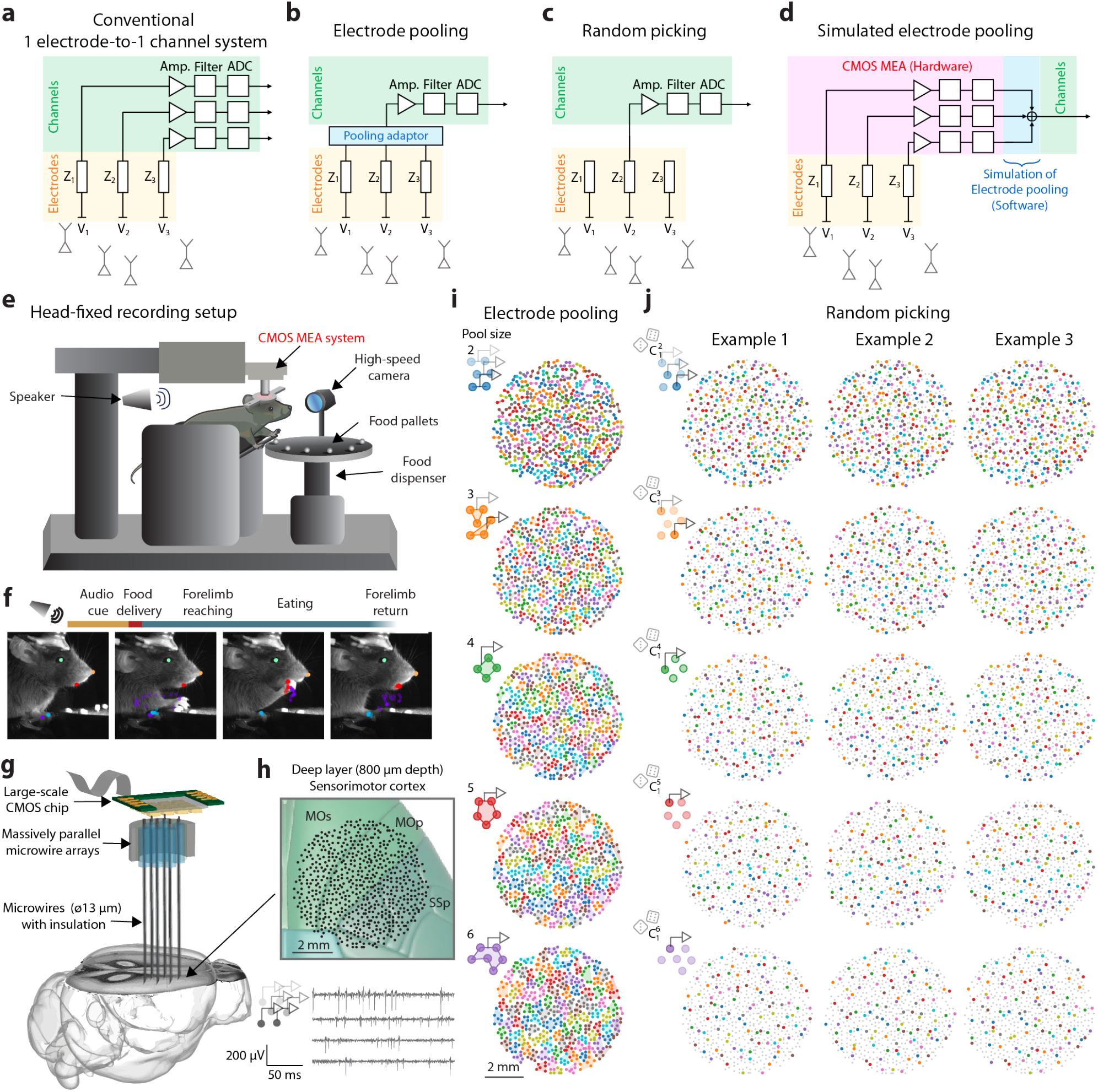
Experimental framework for electrode pooling. **a**, Split mode: each electrode is connected to an independent amplifier, filter and ADC channel. **b**, Electrode pooling: multiple electrodes are tied to a single readout channel through a pooling adaptor. **c**, Random picking control: one electrode from each pool is retained to match the channel count of pooling without signal mixing. **d**, Software implementation used here to emulate hardware pooling by combining split-mode signals acquired through the CMOS MEA system. **e**, Schematic of the cue-guided reach-to-grasp task with simultaneous neural recording and high-speed video tracking. **f**, Example video frames illustrating task epochs and DeepLabCut-tracked forelimb landmarks. **g**, Schematic of the high-density microwire array interfaced to cortex. **h**, *Upper*, representative microwire-array image on the CMOS MEA showing the spatial distribution of implanted microwires across deep-layer sensorimotor cortical regions (800 μm depth), including secondary motor (MOs), primary motor (MOp), and primary somatosensory (SSp) areas. *Lower*, representative example extracellular recording traces. **i**, Electrode maps for pooled configurations (M = 2-6 electrodes per channel). Colored polygons indicate electrodes assigned to the same pooled channel. **j**, Three examples of channel-matched random picking maps for the corresponding pool sizes. The unconnected electrodes are indicated in gray. Scale bars, 2 mm in **h-j**.

Neural activity was recorded with a high-density microwire array interfaced to a CMOS MEA system (**Fig. 1g,h**), and pools of size M = 2-6 were tiled across the array (**Fig. 1i,j**).

All analyses were performed on a single high-density recording dataset from one mouse implanted with one 638-wire array. This dataset contained 180 behavioral trials recorded over approximately 90 min, with approximately 70% successful reach-and-grasp trials. The split-mode recording yielded 2112 curated sorted units. This single-dataset design allowed a direct within-recording comparison of split mode, electrode pooling and channel-matched random picking while eliminating variability in animal identity, implant placement, cortical sampling, behavioral task and recording state.

Having established these modes, we next asked whether pooled recordings could support kinematic decoding. Using population firing rates from sorted units, we trained recurrent neural network (RNN) decoders to predict x- and y-axis paw position (**Fig. 2a**). Split-mode recordings achieved high decoding accuracy (*R*^2^ = 0.94 ± 0.02, mean ± s.d., n = 25 model evaluations). Pooling preserved much of this performance across all pool sizes (M): M = 2, 0.91 ± 0.03; M = 3, 0.88 ± 0.02; M = 4, 0.87 ± 0.02; M = 5, 0.83 ± 0.03; and M = 6, 0.80 ± 0.03 (**Fig. 2b** and **Table 1**). The distribution widened only modestly with pool size, remaining tighter than the corresponding random picking control (s.d. = 0.02-0.04; **Table 1**).

**Fig. 2.**
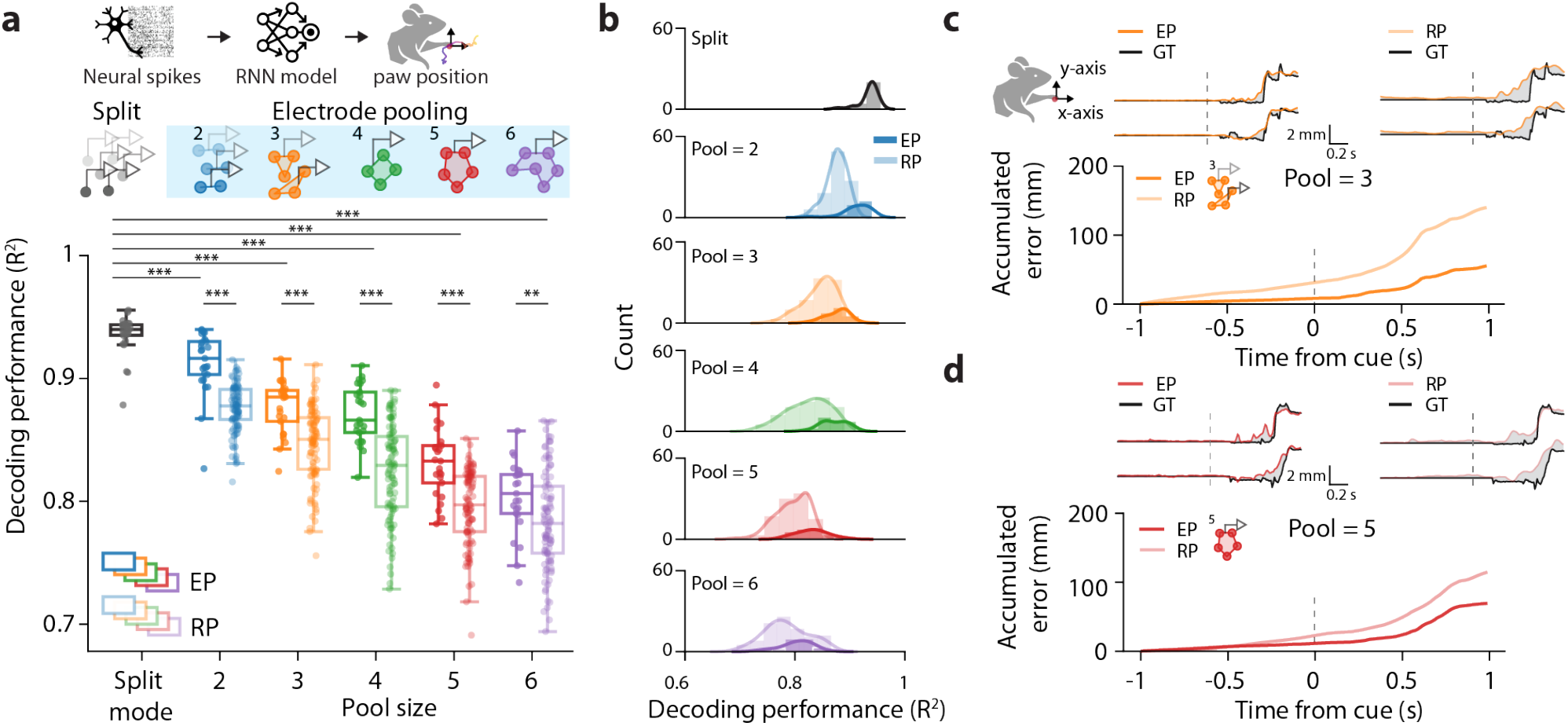
Electrode pooling preserves kinematic decoding better than a channel-matched control. **a**, Decoding workflow and summary of decoding performance. Population firing rates from sorted spikes were used to train a recurrent neural network (RNN) decoder to predict forelimb x- and y-axis position. Box plots show decoding performance (*R*^2^) for split mode, electrode pooling (EP), and random picking (RP). For split mode and EP, n = 25 model evaluations per configuration (fivefold cross-validation × five random initializations). For RP, n = 100 model evaluations per pool size (20 picking configurations × fivefold cross-validation). Boxes show the median and interquartile range; center lines, medians; whiskers, 1.5 times the interquartile range; dots, individual evaluations. Asterisks indicate two-tailed Brunner-Munzel test results for the indicated comparisons (*P < 0.05, **P < 0.01, and ***P < 0.001). **b**, Distributions of *R*^2^ for split mode, EP, and RP. P values for split versus EP and EP versus RP comparisons are listed in **Table 1. c**,**d**, Representative decoded and ground-truth (GT) trajectories and accumulated decoding error for pool sizes M = 3 (**c**) and M = 5 (**d**). The examples shown correspond to median decoding performance for each pool size. EP traces are shown at left and RP traces at right. Gray shading indicates the decoding error relative to the ground-truth trajectory. Dashed vertical lines mark cue onset.

**Table 1.** Statistical comparisons for decoding performance in Fig. 2 Values are mean ± s.d. P values were obtained with two-sided Brunner-Munzel tests.

| Pool size (M) | Split Mode<br>($R^2$, n = 25) | Electrode Pooling<br>($R^2$, n = 25) | Split vs EP<br>P value | Random Picking<br>($R^2$, n = 100) | EP vs RP<br>P value |
| --- | --- | --- | --- | --- | --- |
| Split | 0.94 ± 0.02 | | | | |
| 2 | 0.94 ± 0.02 | 0.91 ± 0.03 | <0.0001 | 0.88 ± 0.02 | <0.0001 |
| 3 | 0.94 ± 0.02 | 0.88 ± 0.02 | <0.0001 | 0.85 ± 0.03 | <0.0001 |
| 4 | 0.94 ± 0.02 | 0.87 ± 0.02 | <0.0001 | 0.82 ± 0.04 | <0.0001 |
| 5 | 0.94 ± 0.02 | 0.83 ± 0.03 | <0.0001 | 0.80 ± 0.03 | <0.0001 |
| 6 | 0.94 ± 0.02 | 0.80 ± 0.03 | <0.0001 | 0.79 ± 0.04 | 0.0085 |

Compared with split mode, pooled decoding was significantly lower at all pool sizes (all two-sided Brunner-Munzel P < 0.0001; **Table 1**). Crucially, electrode pooling outperformed the channel-matched random picking control for all pool sizes tested (M = 2-5, P < 0.0001; M = 6, P = 0.008; **Table 1**). Representative decoded trajectories illustrate that pooled recordings tracked reach kinematics more faithfully than random picking, particularly after movement onset and during the return phase (**Fig. 2c,d**). Thus, pooling compresses channel count more efficiently than simply discarding electrodes.

Because larger pools necessarily use fewer readout channels, raw decoding performance shifted downward as channel count decreased with increasing pool size (**Supplementary Fig. 1a**). We therefore tested whether the gap between electrode pooling and random picking could be explained simply by this loss of channels. After mean-centering decoding performance within each pool size, thereby removing the average effect of reduced channel number, dispersion was broadly similar across compressed channel counts, with split mode showing the narrowest spread and M = 2 slightly more stable than larger pools (**Supplementary Fig. 1b**). When performance was further mean-centered within each specific electrode-selection or pooling configuration, electrode pooling and random picking showed comparable cross-validation- and decoder-initialization-related spread within a given pool size (**Supplementary Fig. 1c**). Thus, the difference between electrode pooling and random picking is not explained by the smaller total channel number at larger pool sizes; instead, the broader random picking distributions arise mainly from which electrode is retained from each pool, reinforcing random picking as an appropriate control for channel reduction without signal mixing.

### Pooling increases informative spike and unit yield per channel

To understand why pooled decoding remained strong, we next examined spike and unit yield (**Fig. 3** and **Table 2**). The random-picking control required substantial independent spike-sorting effort. Because each random-picking configuration retained a different subset of electrodes, its spike population could not be inferred by simply removing units from the split-mode dataset. We therefore repeated spike sorting and manual curation independently for every random-picking configuration. Across M = 2–6, this amounted to 100 independent random-picking spike-sorting analyses, in addition to the split-mode and electrode-pooling datasets. Each configuration contained hundreds to more than one thousand curated units, making the random-picking benchmark a full raw-voltage control rather than a lightweight post hoc subsampling analysis. Pooling mixes signals from multiple electrodes, so the total number of sortable spikes is expected to fall as individual spike amplitudes are diluted and collisions become more frequent. Consistent with this expectation, total spike counts declined monotonically with pool size (**Fig. 3a**) and total unit counts also decreased relative to split mode (**Fig. 3d**). Summary spike and unit yield values are provided in **Table 2**. Yet these totals do not capture the relevant hardware trade-off, because pooling uses fewer channels.

**Fig. 3.**
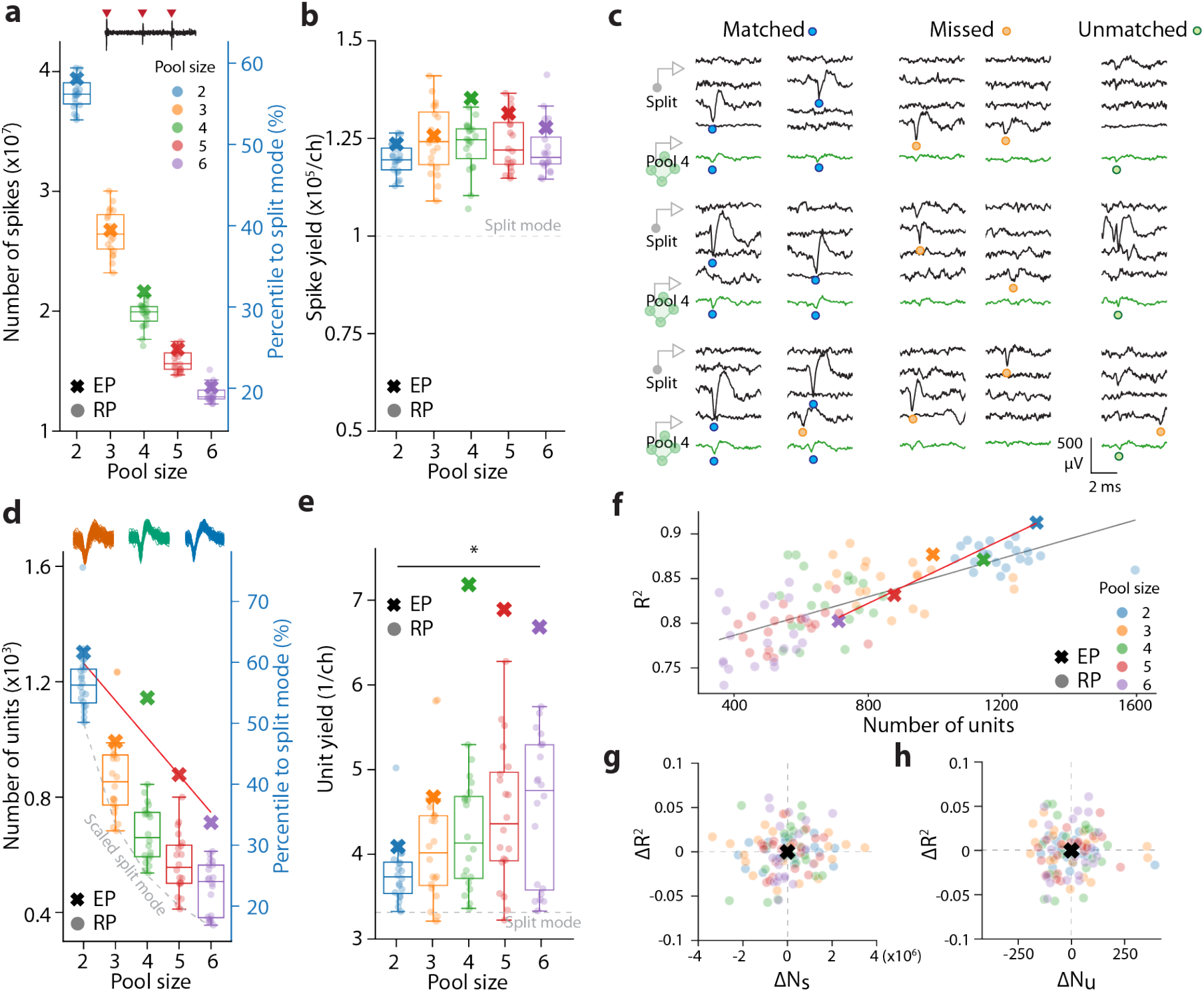
Spike and unit yield after pooling and their relationship to decoding. **a**, Total spike counts across pool sizes for EP and RP. The right axis shows percentile relative to split mode; colored crosses and light circles denote the EP and RP, respectively. **b**, Spike yield per recording channel for EP and RP across pool sizes; the dashed line indicates split-mode yield per channel. **c**, Representative waveform snippets from split-mode recordings and the corresponding pooled recordings for matched, missed, and unmatched spikes (illustrated for M = 4); colored circles denote the timing of the negative peak for each spike. **d**, Total isolated unit counts across pool sizes for EP and RP. The right axis shows percentile relative to split mode; colored crosses and gray circles denote the EP and RP, respectively. Solid line represents the linear regression fit of the EP. Dashed line represents the theoretical lower bound of the number of units identified in EP, calculated by dividing the total isolated unit counts identified in the split mode by the pool size. **e**, Unit yield per channel for EP and RP across pool sizes; the dashed line indicates split-mode yield per channel. **f**, Decoding performance as a function of total unit count across all datasets. Colors denote pool size; colored crosses and gray circles indicate the EP and RP, respectively, for each pool size. Red and grey lines represent the linear regression fit of the EP and RP, respectively. **g**,**h**, Residual analyses relating differences in spike counts (ΔNs) or unit counts (ΔNu) to residual differences in decoding performance (Δ*R*^2^). For each pool size, we first calculated the mean decoding performance, mean spike counts, and mean unit counts across the 20 RP configurations. The RP data were then centered by subtracting these means from the corresponding values within each pool size, thereby isolating relative differences. Because EP includes only a single configuration per pool size, its values were centered at the origin for each pool size. Each dot represents one configuration; black crosses mark EP values. In panels b and e, boxes show the median and interquartile range; center lines, medians; whiskers, 1.5 times the interquartile range; dots, individual configurations. Summary values are listed in **Table 2**. EP, electrode pooling. RP, random picking. Summary values for spike and unit yield are listed in **Table 2**.

**Table 2a.** Spike statistics for Fig. 3. **Split-mode reference (M = 1):** *N*_*S*_=6.8×10^7^, *Y*_*S*_=1.1×10^5^

| Pool size (M) | EP $N_S$ (% of split) | Random Pick $N_S$ (% of split) | EP $Y_S$ | Random Pick $Y_S$ |
| --- | --- | --- | --- | --- |
| 2 | $3.9 \times 10^7$ (58.1%) | $3.8 \pm 0.3 \times 10^7$ ($56.2 \pm 0.4\%$) | $1.24 \times 10^5$ | $1.20 \pm 0.01 \times 10^5$ |
| 3 | $2.7 \times 10^7$ (39.5%) | $2.7 \pm 0.4 \times 10^7$ ($39.2 \pm 0.6\%$) | $1.27 \times 10^5$ | $1.25 \pm 0.02 \times 10^5$ |
| 4 | $2.2 \times 10^7$ (31.9%) | $2.0 \pm 0.2 \times 10^7$ ($29.1 \pm 0.4\%$) | $1.35 \times 10^5$ | $1.23 \pm 0.01 \times 10^5$ |
| 5 | $1.7 \times 10^7$ (24.8%) | $1.6 \pm 0.2 \times 10^7$ ($23.4 \pm 0.3\%$) | $1.31 \times 10^5$ | $1.24 \pm 0.02 \times 10^5$ |
| 6 | $1.4 \times 10^7$ (20.2%) | $1.3 \pm 0.2 \times 10^7$ ($19.3 \pm 0.2\%$) | $1.28 \times 10^5$ | $1.22 \pm 0.02 \times 10^5$ |

**Table 2b.** Unit statistics for Fig. 3. **Split-mode reference (M = 1):** *N*_*U*_=2112, *Y*_*U*_=3.3

| Pool size (M) | EP $N_U$ (% of split) | Random Pick $N_U$ (% of split) | EP $Y_U$ | Random Pick $Y_U$ |
| --- | --- | --- | --- | --- |
| 2 | 1302 (61.7%) | $1202.90 \pm 26.17$ ($57 \pm 1.2\%$) | 4.08 | $3.77 \pm 0.08$ |
| 3 | 993 (47%) | $876.00 \pm 34.56$ ($41.5 \pm 1.6\%$) | 4.66 | $4.11 \pm 0.16$ |
| 4 | 1144 (54.2%) | $674.15 \pm 21.74$ ($31.9 \pm 1\%$) | 7.15 | $4.21 \pm 0.14$ |
| 5 | 878 (41.6%) | $569.05 \pm 23.46$ ($26.9 \pm 1.1\%$) | 6.86 | $4.45 \pm 0.18$ |
| 6 | 712 (33.7%) | $483.85 \pm 20.41$ ($22.9 \pm 1\%$) | 6.65 | $4.52 \pm 0.19$ |

When normalized by acquisition channel, electrode pooling consistently delivered more spikes per channel than random picking across all pool sizes and stayed above the split-mode baseline for intermediate pool sizes (**Fig. 3b** and **Table 2**). Unit yield per channel showed a similar pattern (**Fig. 3e** and **Table 2**): although the absolute number of sorted units fell with increasing M, pooled recordings retained more decodable units per available channel than the channel-matched control. Example waveforms illustrate that many spikes remain recoverable after pooling, alongside expected missed and pooled-only spikes (**Fig. 3c**).

Across acquisition configurations, decoding accuracy scaled positively with total unit count (**Fig. 3f**), indicating that richer sampled populations generally support better kinematic prediction. However, residual analyses showed that the pooled advantage could not be explained solely by changes in total spike or unit numbers (**Fig. 3g,h**). These results suggest that pooling improves not only the quantity of spikes and units per channel, but also the quality of the population information retained within a fixed readout budget.

### Matched units dominate pooled decoding after pooling

Because yield alone did not explain decoding robustness, we next asked how pooling reshaped the sorted population (**Fig. 4**). Pooled units were matched to split-mode units using cosine similarity of spike templates (threshold 0.9; Methods). Pooling preserved a substantial set of matched units across all pool sizes while also generating unmatched (pooled-only) units that were isolated only in the pooled recordings and missed units that were present only in split mode (**Fig. 4a,b**). Spatial maps showed that matched units were broadly distributed across the array, whereas unmatched and missed units were regionally enriched, consistent with local differences in crowding and signal overlap (**Fig. 4c,d**). Representative templates confirmed that matched units retained recognizable waveform structure after pooling, while unmatched units reflected pooled combinations not cleanly recoverable in split mode (**Fig. 4e**).

**Fig. 4.**
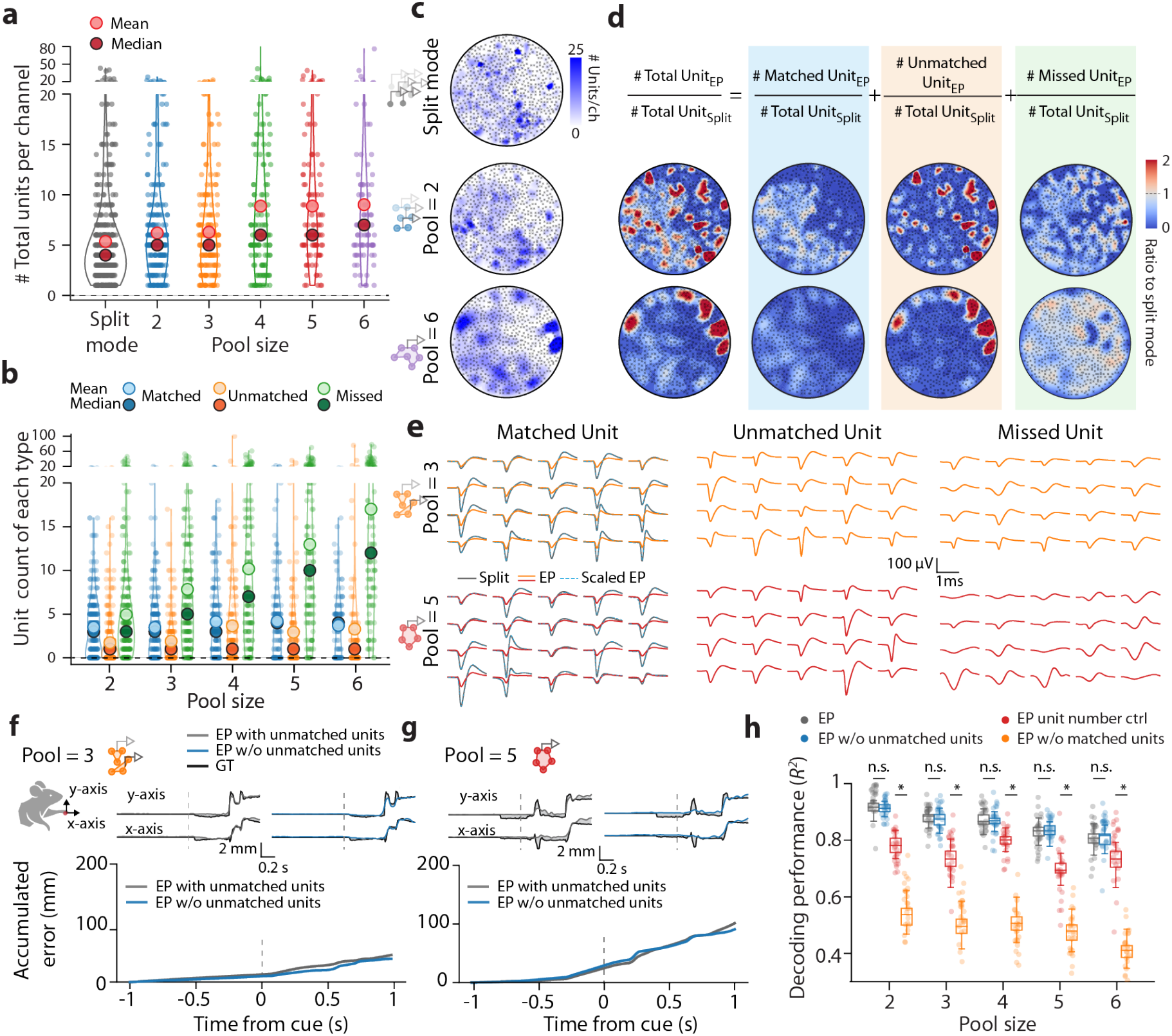
Contribution of matched and unmatched units to decoding. **a**, Distributions of the total number of units per channel for split mode and EP (M = 2-6). Large open and filled circles indicate the mean and median, respectively. **b**, Channel-wise counts of matched, unmatched (pooled-only), and missed units across pool sizes; open and filled circles indicate the mean and median, respectively. **c**, Spatial maps for split mode and representative pooled configurations (M = 2 and M = 6) showing local changes in unit count relative to split mode. **d**, Ratio maps summarizing matched, unmatched, and missed units relative to split mode for representative pooled configurations (M = 2 and M = 6). The color scale indicates the ratio to split mode. Additional configurations are shown in **Supplementary Fig. 2. e**, Example spike templates from matched, unmatched, and missed units for representative pool sizes (M = 3 and M = 5). **f**,**g**, Representative decoded trajectories and cumulative error after removing unmatched units from pooled datasets for pool sizes M = 3 (f) and M = 5 (g); gray, original EP decoding; blue, decoding after removal. **h**, Decoding performance for the original EP datasets, after removing unmatched (pooled-only) units, after removing matched units, or under the unit-number control. In the unit-number control, unmatched units were removed first, followed by a randomly selected subset of matched units, such that the remaining number of matched units equaled the number of unmatched units. Boxes show the median and interquartile range; center lines, medians; whiskers, 1.5 times the interquartile range; dots, individual model evaluations. Statistical comparisons for panel h are listed in **Table 3**. GT, ground truth. EP, electrode pooling.

We then assessed which unit classes contributed to decoding. Removing unmatched (pooled-only) units did not significantly impair decoding performance, particularly at larger pool sizes, whereas removing matched units produced a larger drop across all pool sizes (two-way ANOVA followed by Tukey’s post hoc test with Bonferroni correction; **Fig. 4f-h** and **Table 3**). A unit-number control, in which random units were randomly downsampled to the same remaining count, produced a larger penalty than removing unmatched units alone. Thus, pooled recordings derive most of their decodable kinematic information from units corresponding to split-mode units, while the retention of population-level structure appears more important than the emergence of pooled-only units.

**Table 3.** Pairwise statistics for unit-removal decoding analysis in Fig. 4h. Entries are Bonferroni-adjusted P values from Tukey’s post hoc comparisons after two-way ANOVA. For each configuration, n = 25 model evaluations. EP, original electrode pooling; EP_UM_, electrode pooling after removing unmatched pooled-only units; EP_M_, electrode pooling after removing matched units; Unit Count Ctrl, unit-count-matched control.

| Comparison | M = 2 | M = 3 | M = 4 | M = 5 | M = 6 |
| --- | --- | --- | --- | --- | --- |
| EP vs EP<sub>UM</sub> | 0.9998 | 0.9812 | 0.9445 | 0.9984 | 1.0000 |
| **EP VS CTRL** | <0.0001 | <0.0001 | <0.0001 | <0.0001 | <0.0001 |
| **EP VS EP<sub>M</sub>** | <0.0001 | <0.0001 | <0.0001 | <0.0001 | <0.0001 |
| **EP<sub>UM</sub> VS CTRL** | <0.0001 | <0.0001 | <0.0001 | <0.0001 | <0.0001 |
| **EP<sub>UM</sub> VS EP<sub>M</sub>** | <0.0001 | <0.0001 | <0.0001 | <0.0001 | <0.0001 |
| **CTRL VS EP<sub>M</sub>** | <0.0001 | <0.0001 | <0.0001 | <0.0001 | <0.0001 |

### Pooling preserves low-dimensional latent dynamics of population activity

This unit-level analysis pointed to a population-level explanation for pooling performance: decoding may remain robust if pooling preserves the low-dimensional activity patterns that drive movement. We therefore inferred single-trial latent factors using latent factor analysis via dynamical systems (LFADS)^31^ and visualized peri-event population patterns around cue onset and movement execution (**Fig. 5a-d**). Across pool sizes, electrode-pooling peri-event histograms retained structured temporal modulation that closely resembled split mode, whereas random picking produced more variable and often attenuated patterns depending on the selected configuration.

**Fig. 5.**
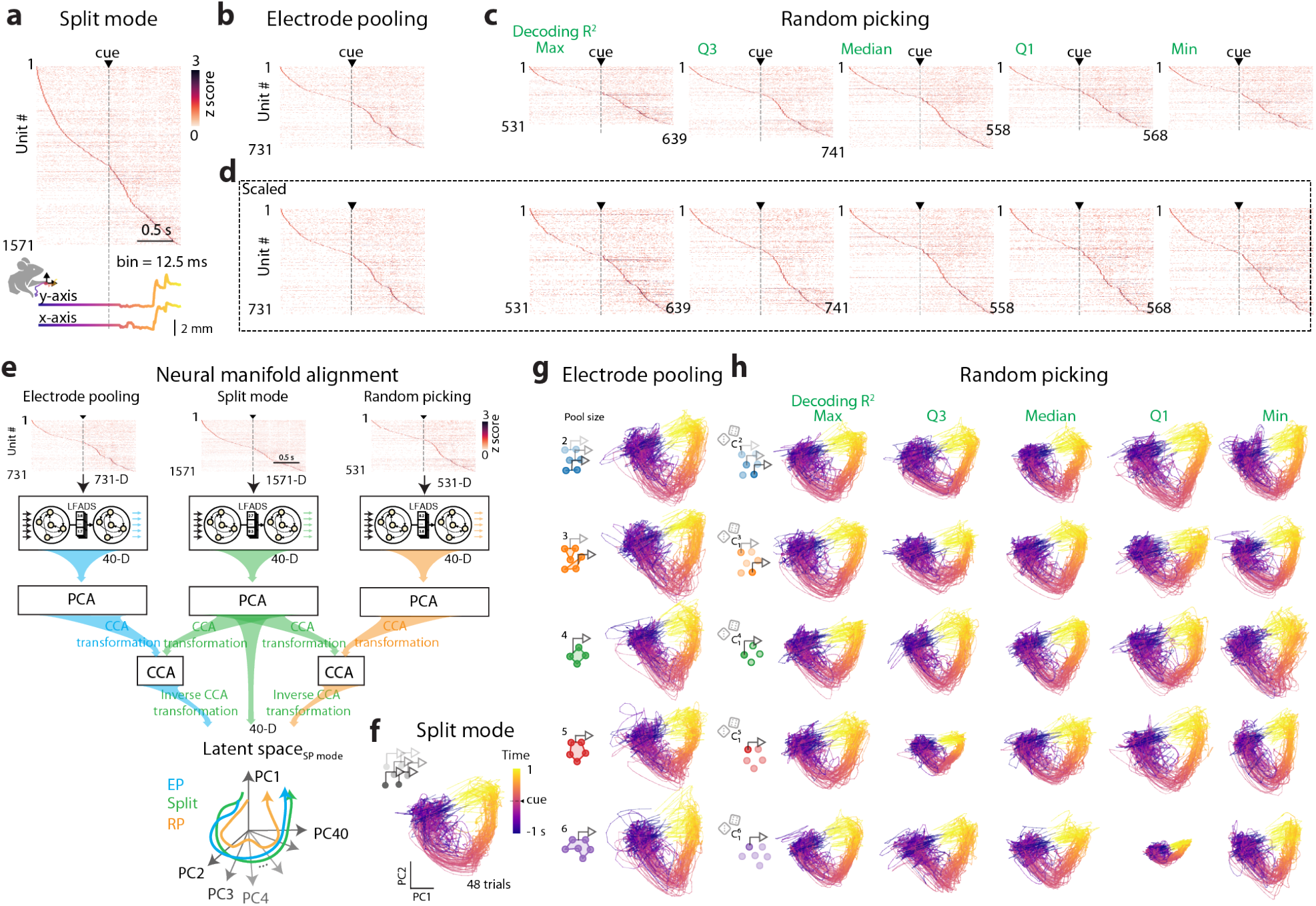
Electrode pooling preserves latent population trajectories. **a**, Split-mode peri-event time histograms (PETHs) for sorted units aligned to cue onset and ordered by peak activity. **b**, PETHs from representative pooled recordings (M = 3). **c**, PETHs from random picking (RP) configurations spanning the distribution of decoding performance (maximum, third quartile, median, first quartile, and minimum). **d**, Scaled PETHs of both electrode pooling and random picking with the horizontal axis normalized to the same height across both methods. These scaled PETHs are used to evaluate the similarity of neuronal firing patterns across modes. **e**, Neural manifold alignment. Latent factors (40 dimensions) were inferred from firing rates with LFADS, capturing latent dynamics. Principal component analysis was applied to trial-averaged latent factors from split mode, EP, and RP, using data from 48 successful reach trials spanning 1 s before to 1 s after cue onset. For cross-configuration comparison, canonical correlation analysis was used to align split-mode principal component scores with those from EP and RP (see Methods, “Latent-dynamics analysis”; **Supplementary Fig. 4**). **f**, Representative split-mode smoothed latent trajectories colored by time and overlapping 48 trials. **g**,**h**, Aligned latent trajectories for EP (**g**) and RP (**h**) across pool sizes and selected RP quantiles. The number of recording channels for both electrode pooling and random picking are identical in each row (48 trials). Color indicates time from cue onset. Scale bars, 0.5 s in **a-d**. EP, electrode pooling. RP, random picking.

To compare latent trajectories directly across acquisition modes, we aligned pooled and control latent activity to the split-mode principal-component space using canonical correlation analysis (**Fig. 5e,f**). Electrode-pooling latent trajectories consistently overlapped the split-mode latent trajectories across pool sizes. In contrast, latent trajectories derived from random picking exhibited greater variability, dispersing broadly across best-to worst-decoding configurations (**Fig. 5g,h**). These qualitative differences suggest that pooling preferentially preserves the dominant population geometry linked to behavior.

### Latent-dynamics similarity explains pooled decoding robustness

We next quantified the latent-dynamics preservation suggested by **Fig. 5**. Projecting pooled activity into the split-mode reference principal-component space recovered the major split-mode temporal structure for representative pool sizes of 3 and 5 (**Fig. 6a,b**). Random picking could approximate the best-performing pooled configurations but showed much larger variability across the distribution of control configurations (**Fig. 6c**).

**Fig. 6.**
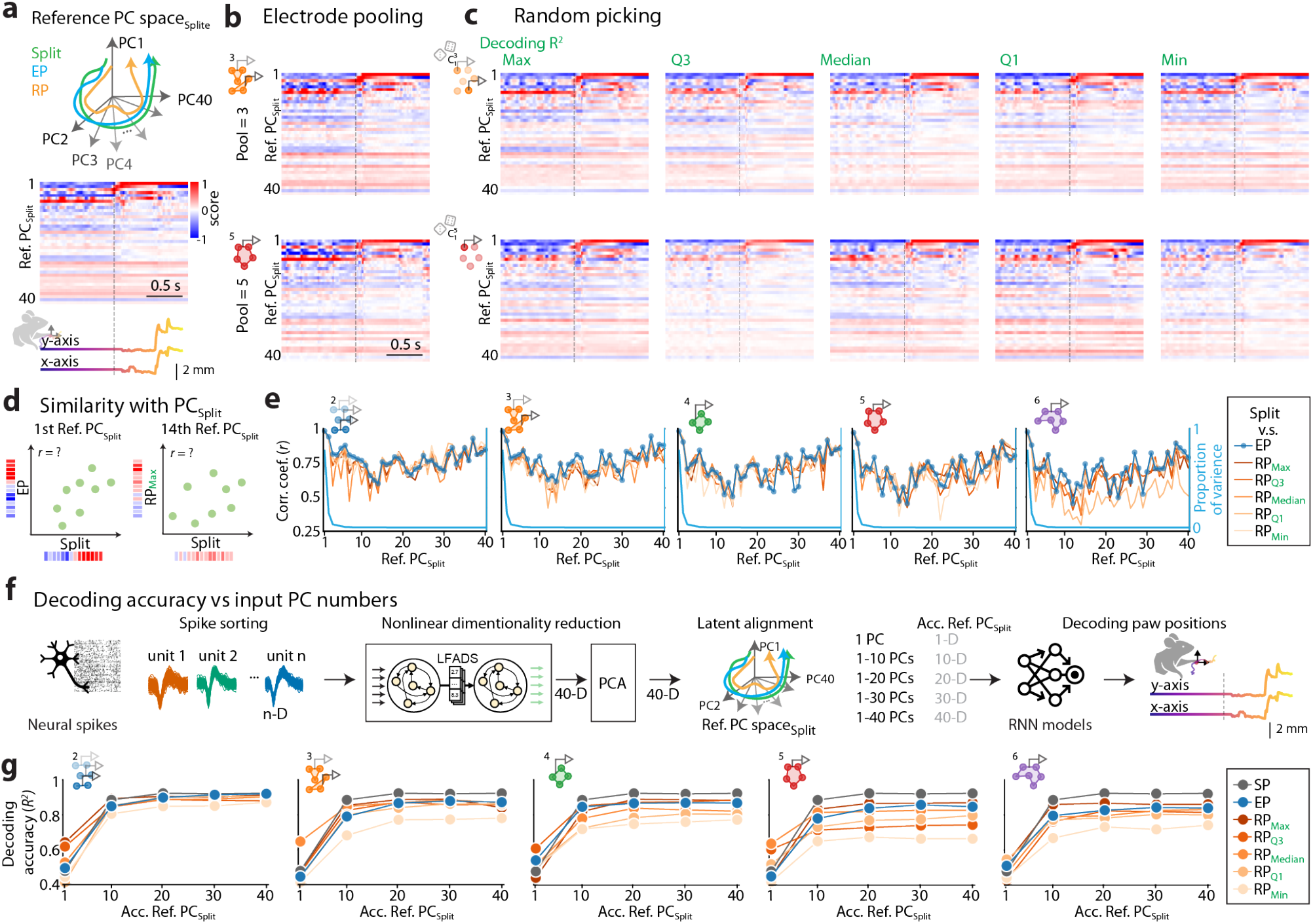
Quantitative comparison of pooled and control latent dynamics. **a**, Split-mode reference principal-component (Ref. PCSplit) space used for alignment. The heat map below shows reference split-mode PC scores. The bottom panel presents the corresponding decoded kinematics derived from these scores. **b**, EP latent activity projected into the split-mode reference PC space for representative pool sizes (M = 3 and M = 5). **c**, RP projections from configurations spanning the decoding-performance distribution. **d**, Schematic for quantifying similarity, defined by correlation coefficients calculated independently for each PC, between aligned EP or RP PC scores and the split-mode reference PC scores. **e**, Similarity as a function of reference PC number for EP and RP across pool sizes. The blue line indicates the proportion of split-mode variance explained by the reference PCs. **f**, Schematic illustrating the evaluation of input PC numbers on decoding performance. Accumulated latent activity projected into the split-mode reference PC space (Acc. Ref. PCSplit) was used as decoder input. **g**, Decoding performance obtained with Acc. Ref. PCSplit when progressively increasing numbers of PCs from each configuration (split mode, aligned EP, or aligned RP) were used as inputs. SP, split mode; EP, electrode pooling; RP, random picking.

Similarity to split mode remained highest in the leading reference principal components and declined gradually across higher dimensions for electrode pooling, reflecting consistent recovery of structure. In contrast, random picking lost correspondence more rapidly and less consistently (**Fig. 6d,e**). Using the aligned principal components as progressively larger decoder inputs (**Fig. 6f**), pooled activity retained more task-relevant low-dimensional information than random picking across all pool sizes (**Fig. 6g**). These results connect pooled decoding robustness to preservation of the latent manifold rather than perfect recovery of every individual spike train.

### Additional electrodes per channel improve decoding under a fixed channel budget

Finally, we tested the design implication of these findings: pooling should not only reduce the readout channels needed for an existing array, but also allow broader spatial sampling when channel count is fixed. We therefore fixed the available readout budget to 107 channels and varied how many electrodes were connected to each channel, corresponding to a progressively larger electrode inventory (**Fig. 7a-c**).

**Fig. 7.**
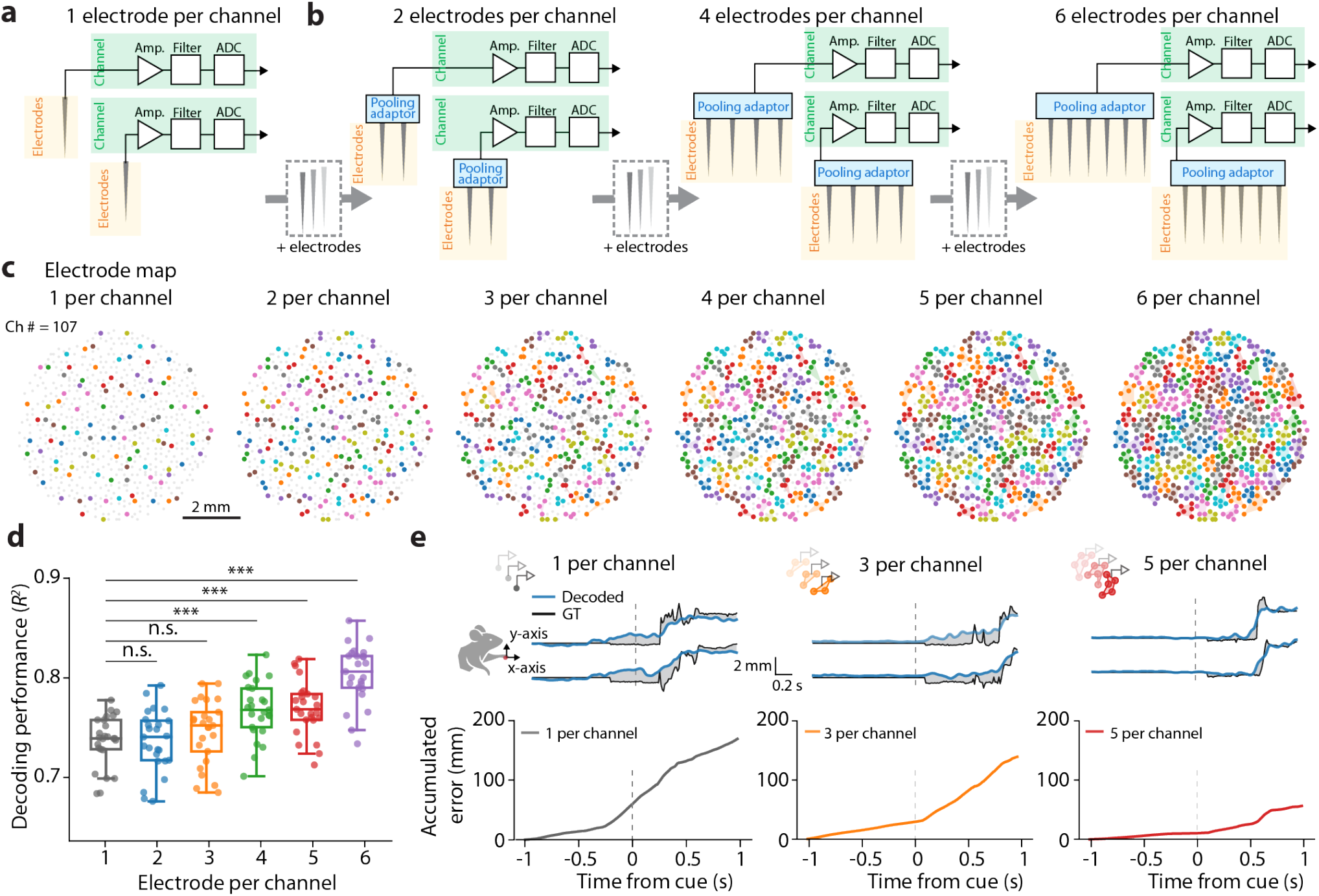
More electrodes per channel improve decoding under a fixed channel budget. **a**,**b**, Schematic of scaling from one electrode per channel to pooled acquisition with 2, 4, or 6 electrodes per channel while keeping the readout budget fixed. **c**, Corresponding electrode maps for a fixed channel budget of 107 channels. Colored polygons indicate electrodes combined on the same pooled channel. **d**, Decoding performance as a function of electrodes per channel. Boxes show the median and interquartile range; center lines, medians; whiskers, 1.5 times the interquartile range; dots, individual model evaluations. n.s., not significant; ***P < 0.001 by the statistical test described in Methods. **e**, Representative decoded and ground-truth trajectories (top) and accumulated error (bottom) for one, three, and five electrodes per channel. Dashed vertical lines mark cue onset.

Under this constraint, decoding improved as more electrodes were attached to each channel (**Fig. 7d**). Performance was similar between one and three electrodes per channel, but increased significantly for four to six electrodes per channel, with the highest accuracy obtained at six electrodes per channel. Representative decoded trajectories and cumulative decoding errors illustrate the same trend: broader spatial sampling reduced trajectory error even though each individual channel carried mixed signals (**Fig. 7e**). Pooling therefore turns unused spatial capacity into behaviorally informative population coverage.

## Discussion

This study shows that electrode pooling can substantially relax the readout bottleneck of large-scale extracellular recording without a proportional loss in decoding performance. Even with 4-6 electrodes combined on each channel, pooled recordings retained robust prediction of forelimb kinematics and outperformed a channel-matched strategy that simply discarded electrodes. In other words, the dominant cost of pooling is not equivalent to losing three to five of every four to six electrodes; instead, mixed signals still carry recoverable, behaviorally useful population information.

The results also clarify why pooling works. Pooling reduced total spike and unit counts, as expected from amplitude averaging and spike overlap, yet preserved the low-dimensional latent dynamics most tightly linked to behavior. This observation resonates with a growing view that population-level neural computation and decoding depend more on stable latent manifolds than on the perfect identity of individual units^21–23,27,28,30^. By retaining those shared dynamics, pooled recordings remain decodable even when some single units are distorted or lost.

Our matched-unit analysis reinforces this interpretation. Unmatched (pooled-only) units appeared after pooling, but targeted unit-removal analyses indicated that the pooled units corresponding to split-mode units carried most of the decodable kinematic information. The practical implication is that pooling should not be judged solely by traditional single-unit recovery metrics. For applications such as neural decoding, closed-loop control or latent-state estimation, the more relevant question is how much task-relevant population geometry survives channel compression. Our data indicate that pooling preserves task-relevant neural information far beyond what would be expected from the reduction in channel count alone.

Electrode pooling is also attractive because it complements, rather than replaces, ongoing advances in dense neural hardware. Switchable probes, multiplexed arrays, CMOS-integrated systems, ultrasmall bioactive microelectrodes, flexible kirigami microelectrode arrays and quantitative low-trauma probe designs continue to expand electrode density and reduce implantation burden^2–4,7–10,15,18,19^, but channel budget, power, routing and packaging remain limiting. Pooling provides a lightweight architectural option that can trade some per-electrode separability for much broader spatial sampling. Our fixed-budget analysis highlights this advantage directly: when channel count was held constant, adding more electrodes per channel improved decoding.

Several limitations should guide future work. First, the neural recording results were obtained from one mouse, one implanted array and one recording session. The present study should therefore be interpreted as a high-density within-dataset proof-of-concept benchmark of electrode pooling, rather than as a population-level estimate of performance across animals. This design allowed stringent comparison of split mode, electrode pooling and random picking under matched electrode geometry, behavioral state and recording configurations, but systematic multi-animal and multi-session studies will be required to establish generalization across implants, cortical areas, behaviors and chronic recording conditions. Second, the optimal pool geometry is likely to depend on electrode spacing, waveform diversity and circuit statistics; pooling nearby redundant electrodes may differ from pooling more spatially diverse sites. Third, most pooled data here were generated by software simulation on split-mode recordings; dedicated switchable hardware or pooling adaptors will be essential to quantify real-time noise, impedance and chronic stability effects *in vivo*. Finally, future studies should couple pooling with adaptive decoding and online latent-space alignment to determine how far channel compression can be pushed in chronic interfaces. Despite these open questions, our results identify electrode pooling as a practical route to scale neural recordings by preserving the population dynamics that matter most for behavior.

## Methods

### Overview of electrode pooling method

Electrode pooling was implemented to evaluate whether multiple electrodes could be combined into single acquisition channels without loss of behaviorally relevant neural information. Neural activity was recorded using a high-density microwire electrode array interfaced with a 2D CMOS microelectrode array system (**Fig. 1g,h**). In conventional split mode, each electrode was assigned to a distinct acquisition channel. In pooled mode, multiple electrodes were assigned to a single channel. To perform software-based pooling, raw voltage traces from electrodes within a pool were averaged to generate a composite signal (**Fig. 1d**). As a control, a random-picking configuration was used to match the channel count of electrode pooling. In this configuration, one electrode was randomly selected from each pool, and its split-mode signal was designated as the channel output (**Fig. 1c,j**).

To assess the impact of electrode pooling on neural information content, we employed three complementary analyses: neural decoding, unit-level recovery, and preservation of latent neural dynamics across split mode, electrode pooling, and random picking. Neural decoding was performed using a recurrent neural network (RNN) designed to predict forelimb kinematics from population firing rates. Decoder performance was quantified by the coefficient of determination between predicted and ground-truth trajectories, averaged across both axes to yield a single metric. Unit-level recovery was assessed with unit matching analysis, which determined whether individual units recorded in split mode could be reliably recovered after pooling. This analysis enabled quantification of units that were preserved, missed, or uniquely detected under pooling.

Latent-dynamics analysis was used to evaluate whether electrode pooling preserved behaviorally relevant low-dimensional neural representations. Latent activity was inferred using LFADS. Canonical correlation analysis (CCA) was applied to align pooled and random-picking trajectories to the split-mode reference space. Similarity between configurations was quantified by correlations of time-resolved trajectories in the aligned space. To further test the retention of task-relevant information, neural decoders were trained on progressively larger sets of principal component scores derived from each configuration.

### Experimental design and ethics

A C57BL/6J mouse (10 weeks old) was used for the reach-and-grasp task. The mouse was housed in groups of 4-5 per cage in a temperature-controlled environment (22 ± 1°C) with a 12-hour light/dark cycle. Food and water were available *ad libitum*. All animal procedures were reviewed and approved by the Institutional Animal Care and Use Committee (IACUC) at Academia Sinica and were performed in accordance with institutional guidelines and applicable national regulations for the scientific use of laboratory animals.

### Experimental sample, recording dataset and trial inclusion

All neural decoding and electrode-pooling analyses were performed on one high-density recording dataset obtained from one mouse implanted with one microwire-CMOS array. The dataset consisted of one recording session lasting approximately 125 min and containing 250 cue-guided reach-and-grasp trials. Each trial consists 25-30s of recording. The implanted array contained 638 usable microwires. To ensure the reliability of kinematic decoding, 181 trials were selected from a total of 250 that met the success criteria defined below. Early trials were excluded due to residual anesthetic effects and insufficient proficiency in food-grasping behavior, whereas later trials were omitted because satiety reduced the mouse’s motivation to attempt grasping. The remaining 181 trials were used for downstream decoding analyses. The split-mode dataset yielded 2112 manually curated sorted units after spike sorting and quality-control filtering. Pooled and random-picking datasets were generated from the same split-mode voltage recordings, allowing direct within-dataset comparison of acquisition strategies while holding constant the animal, implant, cortical region, behavioral task, electrode geometry and recording epoch.

The goal of this study was to benchmark how different readout strategies affected the information retained from the same dense neural recording, rather than to estimate inter-animal biological variability. Accordingly, the primary comparisons among split mode, electrode pooling and random picking were performed within this single high-density dataset. Reported sample sizes for decoder analyses refer to cross-validation folds, decoder initializations and/or independently generated channel configurations, as specified below, and should not be interpreted as independent biological replicates.

Because the neural dataset was obtained from one mouse and one implanted array, statistical comparisons were used to quantify within-dataset differences among readout strategies, channel configurations and decoder evaluations. These tests do not estimate inter-animal variability. For split mode and electrode pooling, n denotes repeated decoder evaluations from cross-validation folds and decoder initializations. For random picking, n denotes evaluations across independently generated channel configurations and cross-validation folds. The biological sample size for the neural recording experiment was N = 1 mouse, N = 1 implanted array and N = 1 recording session.

### Behavioral task and kinematic tracking

The implanted mouse performed a cue-driven, head-fixed reach-and-grasp task (**Fig. 1e,f**). Animals were positioned in a custom behavioral apparatus in front of a pellet-dispensing wheel, with the resting position of the forepaws constrained by a perch to reduce trial-to-trial variability in movement trajectory. The behavioral apparatus was housed in a dark box. Food pellets (20 mg; Bio-Serv, USA) were delivered by rotating the pellet wheel with a servomotor controlled by custom Arduino software approximately 200 ms after the onset of an auditory tone. The cue prompted the animal to reach for the pellet, grasp it, bring it to the mouth, consume it, and return the forelimb to the resting position. Trials were included as successful reach trials when the animal reached the pellet within 1 s after the auditory cue on the first reach attempt and successfully grasped and consumed the pellet. Failed trials were defined as trials in which the animal reached the pellet within 1 s after the auditory cue on the first attempt but failed to grasp and consume the pellet. The downstream electrode-pooling analyses used trial-aligned paw trajectories from successful trials; the latent-dynamics analysis used 48 successful reach trials spanning 1 s before to 1 s after cue onset.

Behavior was recorded with two synchronized high-speed, high-resolution monochrome cameras (Grasshopper 3 GS3-U3-23S6M-C, FLIR, USA) equipped with 6-15 mm (f/1.4) C-mount lenses (Tokina, Japan), positioned perpendicular to one another in front of and to the left of the animal. An infrared LED light source was mounted above each camera. Videos were acquired at 80 frames/s with resolutions of 512 × 344 (front) and 768 × 502 pixels (side). The two cameras were synchronized to each other during acquisition, and behavioral time series were aligned to auditory cue onset.

Forelimb kinematics were extracted using DeepLabCut^32^. For the reaching behavior pipeline, 20 video frames per session were selected by clustering based on visual appearance and manually annotated to identify paw locations. The model was trained using a ResNet-50 backbone with default parameters and subsequently applied to predict paw positions across all video recordings. Automated annotations were visually inspected, and mislabeled frames were manually corrected to ensure accuracy. The resulting trial-aligned paw trajectories served as ground-truth data for downstream analyses of decoding performance under three acquisition configurations: split mode, electrode pooling, and random picking.

### Neural recording and pooling configurations

Neural activity was recorded with a high-density microwire electrode array interfaced to a CMOS voltage-amplifier/readout system described previously (**Fig. 1g,h)**. The system supports simultaneous high-channel-count neural acquisition at 32 kHz per channel with 12-bit digitization. The CMOS sensor uses an AC-coupled low-noise amplifier chain composed of two cascaded common-source gain stages, each with a nominal gain of approximately 10 V/V, yielding a front-end gain of approximately 100 V/V. Peripheral differential-output amplifiers bring the total signal-chain gain to approximately 800 V/V. The amplifier high-pass corner is tunable from approximately 18 to 300 Hz, and the anti-aliasing low-pass filter was configured with 20-kHz corner settings, corresponding to an approximately 12-kHz 3-dB point. The readout electronics include high-speed ADCs and an FPGA for digitization, demultiplexing, packetization and optical data transfer. In the present study, 638 microwires with usable neural signals were analyzed. Raw extracellular voltage traces were acquired at 32 kHz per channel, stored for offline analysis, and digitally band-pass filtered at 300–6000 Hz for spike sorting.

Two acquisition modes were analyzed. In split mode, one electrode was assigned to one acquisition channel. In pooled mode, multiple electrodes were assigned to a single channel. For software electrode pooling, pooled-mode recordings were simulated by averaging the raw voltage traces from all electrodes within the same pool (**Fig. 1d**). Pools of size M = 2-6 were constructed according to the electrode maps shown in **Fig. 1i** using a software pooling algorithm (see **Algorithm 1**). This algorithm comprises two sequential phases: initial spatial partitioning and iterative refinement. In the first phase, the set of electrode coordinates 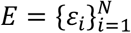 was partitioned into pools of fixed size *M*. The process iteratively identified the electrode furthest from the centroid of *E*. A pool *P*_*k*_ was then formed by ε* and its *M* − 1 nearest ungrouped neighbors. Any remaining electrodes totaling fewer than *M* were grouped into a final residual pool, yielding an initial set of electrode pools 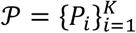. The second phase refined these pools over *T* iterations to enhance spatial compactness by minimizing maximum intra-pool distances. For every unique pair of pools (*P*_*i*_, *P*_*j*_), the algorithm identified electrodes 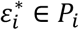 and 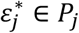 located farthest from their respective centroids. Trial pools 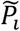 and 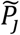 were generated by swapping 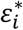 and 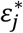. A swap was accepted only if it reduced the pool diameter (diam((*P*)), defined as the maximum distance between any two electrodes within a pool. Through this iterative swap mechanism, the algorithm progressively optimized the spatial compactness of all electrode pools.

#### Algorithm 1

**software electrode pooling**

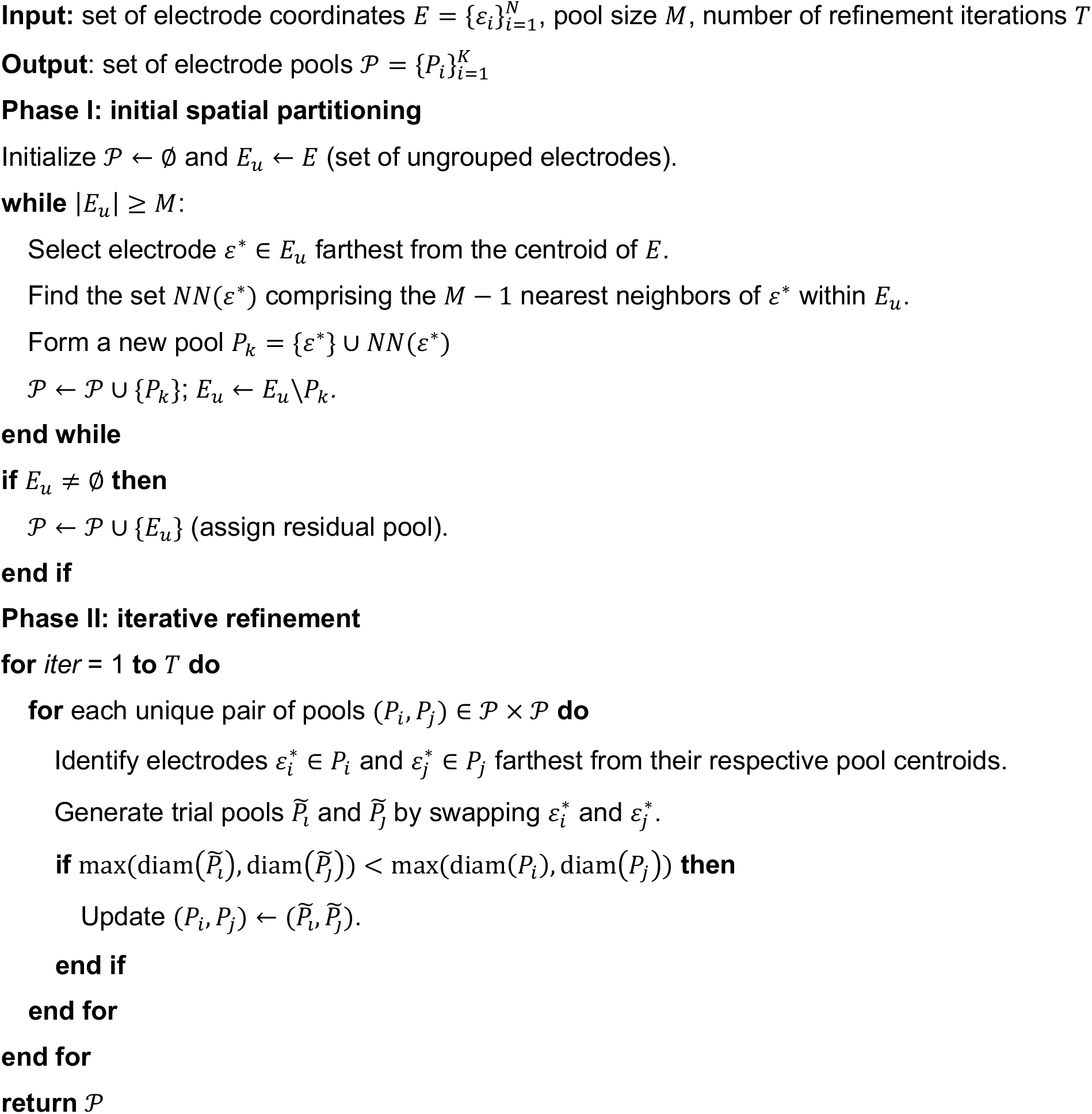

### Random-picking control and independent spike sorting

Random picking served as a channel-matched control for electrode pooling. For each predefined electrode pool, one electrode was randomly selected (See examples in **Fig. 1c,j)** and its split-mode voltage trace was retained as the channel output, while the remaining electrodes in that pool were discarded. This procedure preserved the same output channel count and spatial tiling as electrode pooling, but eliminated signal mixing. For each pool size, 20 independent random-picking configurations were generated.

Importantly, random picking was implemented at the raw-voltage level rather than by *post hoc* deletion of already sorted split-mode units. Each random-picking configuration was therefore spike sorted independently using the same spike-sorting and manual-curation pipeline applied to split mode and electrode pooling. Across the five tested pool sizes, this produced 100 independently spike-sorted random-picking datasets. This design allowed the random-picking control to capture the full effects of electrode selection on spike detectability, waveform separation, unit yield and downstream decoding, rather than only the effects of removing units from a pre-existing sorted population.

### Spike sorting and firing-rate estimation

Spike sorting was performed with KiloSort 2^33^. Sorted units were manually curated to remove units with firing rates below 0.5 Hz, waveforms with positive-to-negative peak-to-peak amplitudes above 3000 μV, and waveforms with negative peaks more positive than −40 μV. The same curation criteria were applied across split mode, electrode pooling, and random picking datasets.

Smoothed firing rates were computed from sorted spike trains using the Elephant analysis framework^34^. Specifically, sorted spike trains for each unit were binned into 12.5-ms time windows, smoothed with a Gaussian kernel (sigma = 1.5 bins), and z-score transformed.

### Neural decoding

Forelimb kinematics were decoded from population firing rates using a recurrent neural network decoder that predicted paw trajectories along the x- and y-axes. Decoder performance was quantified by the coefficient of determination (*R*^2^) between predicted and ground-truth trajectories. To obtain a single performance metric, the *R*^2^ values for the x- and y-axes were averaged with uniform weighting.

The input to the decoder consisted of firing rates from C units recorded across 80 time bins, corresponding to one second of historical neural activity. These inputs were processed by two stacked bidirectional gated recurrent units (BiGRUs), each with a hidden state dimensionality of 1024. The BiGRUs were employed to exploit the sequential structure of firing rates and to capture the temporal dynamics of neural activity. The BiGRU outputs were subsequently passed to a multilayer perceptron (MLP) with 256 hidden units and rectified linear unit activation. The final output of the MLP corresponded to the predicted paw trajectories along the x- and y-axes.

The neural decoder was trained in a supervised manner by minimizing the mean squared error (MSE) loss, defined as

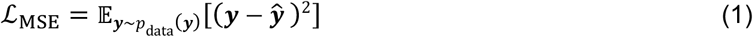

where y and ŷ denote the ground-truth and predicted paw trajectories, respectively, and y ~ p_ data(y) indicates that y is sampled from the data distribution. To mitigate overfitting, dropout regularization^35^ was applied at a rate of 0.05 to each fully connected layer within both the BiGRUs and the MLP. Training was performed using the Adam optimizer^36^ with zero weight decay, employing coefficients of 0.9 and 0.999 for the moving average of gradients and squared gradients, respectively.

Training was subject to an early stopping criterion, whereby learning was terminated if the validation loss failed to improve for ten consecutive epochs. All training procedures were conducted on an Intel Core i7-14700K processor paired with an NVIDIA GeForce RTX 3080 GPU.

For split mode and electrode pooling, decoding performance was evaluated with fivefold cross-validation and five random decoder initializations per fold, yielding n = 25 model evaluations per configuration. For random picking, each of the 20 configurations was evaluated with the same five cross-validation folds but only one random decoder initialization per fold, yielding n = 100 model evaluations per pool size.

For latent analysis, the decoding procedure was analogous to that used for firing rates, with the primary difference being the input representation. Instead of raw firing rates, the decoder received principal component scores derived from neural activity. Specifically, the input consisted of 40-dimensional principal component scores recorded across 80 time bins, corresponding to one second of latent dynamics.

### Unit matching analysis

To assess which split-mode units were recovered after pooling, spike templates from split mode and pooled recordings were compared using cosine similarity. A split-mode unit was labeled as matched if its best pooled-mode match exceeded a similarity threshold of 0.9. Split-mode units without such a match were labeled as missed, and pooled units that were not assigned to any split-mode unit were labeled as unmatched.

### Latent-dynamics analysis

To assess whether electrode pooling preserves behaviorally relevant latent dynamics, we inferred 40-dimensional latent activity from firing rates using LFADS^31^. LFADS is a sequential variational auto-encoder with recurrent neural networks that models latent dynamics underlying observed population activity and is trained to generate “denoised” single-trial firing rates from neural spiking data. The LFADS Run Manager, developed by Daniel J. O’Shea and available on GitHub (https://lfads.github.io/lfads-run-manager), was used to manage training and inference.

The structure of the inferred latent dynamics was examined by applying principal component analysis (PCA) to trial-averaged latent trajectories obtained under three configurations: split mode, electrode pooling, and random picking. Data were drawn from 48 successful reach trials spanning a temporal window from 1 s before to 1 s after cue onset. Projections onto the respective principal component spaces were defined as principal component scores, which matched the dimensionality of the latent activity (40 dimensions).

For comparative analysis, the split-mode space was designated as the reference. Principal component scores derived from electrode pooling and random picking were aligned to this reference space using canonical correlation analysis, as illustrated in **Fig. 5e,f** and **Supplementary Fig. 4**. To enhance visualization, latent dynamics were smoothed using a Savitzky–Golay filter with a window length of 11 and a third-order polynomial.

Similarity of principal component scores between configurations (split-mode, EP, and RP) was quantified using correlations between time-resolved trajectories in the aligned split-mode reference space. To test how much task-relevant information was retained in low-dimensional latent dynamics, neural decoders were trained with progressively larger sets of split-mode, aligned EP, or aligned RP principal component scores (**Fig. 6g**).

### Statistics and reproducibility

All statistical tests were two-sided with a significance threshold of 0.05 unless stated otherwise. Decoding-performance distributions were compared using Brunner-Munzel tests. Spike and unit yields across multiple groups were compared using Kruskal-Wallis tests followed by Dunn’s post hoc multiple-comparison tests with correction for multiple testing. To evaluate decoding performance across groups, a two-way ANOVA was applied, followed by Tukey’s post hoc test with Bonferroni correction. Exact P values for the key decoding comparisons are reported in **Table 1**, spike and unit yield summaries are reported in **Table 2**, and unit-removal comparisons in **Fig. 4** are summarized in **Table 3**.

For split mode and electrode pooling, n = 25 denotes five cross-validation folds with five random decoder initializations per fold. For random picking, n = 100 denotes 20 independently generated channel configurations, each evaluated with the same five cross-validation folds. Box plots show the median, interquartile range, and whiskers extending to 1.5 times the interquartile range unless otherwise noted. Individual data points are plotted throughout.

## Data availability

The datasets generated and analyzed in this study, including raw extracellular recordings, pooled-mode simulations, curated spike times, firing-rate matrices, paw kinematics, decoding outputs, latent variables, and source data underlying **Figs. 2–7** and **Tables 1–3**, will be deposited in Zenodo or an equivalent public repository before publication. During peer review, data are available from the corresponding authors upon reasonable request. Repository names, accession numbers, persistent links, and any access restrictions will be added before final submission.

## Code availability

Custom code used for electrode-pooling simulation, spike-to-rate preprocessing, neural decoding, unit matching, latent-dynamics analysis, and figure generation will be maintained in a versioned GitHub repository and archived with a DOI through Zenodo or an equivalent repository before publication. During peer review, code is available from the corresponding authors upon reasonable request. The final submission will provide the repository URL, release version, and archive DOI.

## Acknowledgements

We sincerely thank Dr. Ching-Lung Hsu, Dr. Yang-Hsien Lin of NVIDIA, Pei-En Wu and members of the Yang and Wu laboratories for helpful discussions. This work was supported by the National Science and Technology Council, Taiwan (NSTC 112-2628-E-006-013 to S.-H.Y.; NSTC 113-2321-B-001-012, 114-2321-B-001-005, NSTC 113-2320-B-001-023 to Y.-W.W), and Academia Sinica (AS-CDA-110-L05 and AS-IAIA-114-L01 to Y.-W.W.). We also thank the National Center for High-performance Computing (NCHC) and NVIDIA-NCKU Artificial Intelligence and High-Performance Computing Application Research Program for providing computational and storage resources.

## Author contributions

S.-H.Y. and Y.-W.W. conceived the study. S.-H.Y. and Y.-W.W. designed the experiments and analysis framework. S.-H.Y., Y.-S.L., Y.-K.C., and Y.-W.W. established the analysis pipeline and graphical user interface. Y.-C.L., W.-Y.H., Y.-F.C., W.-J.C., Y.-S.L., Y.-K.C., Y.-T. C., S.-S.S., S.-H.Y. and Y.-W.W. analyzed the data, prepared the figures and wrote the manuscript with input from all authors. All authors discussed the results. S.-H.Y. and Y.-W.W. wrote the manuscript with contribution of all coauthors.

