## Supplementary Figures for "Electrode pooling preserves movement decoding by retaining neural population dynamics"

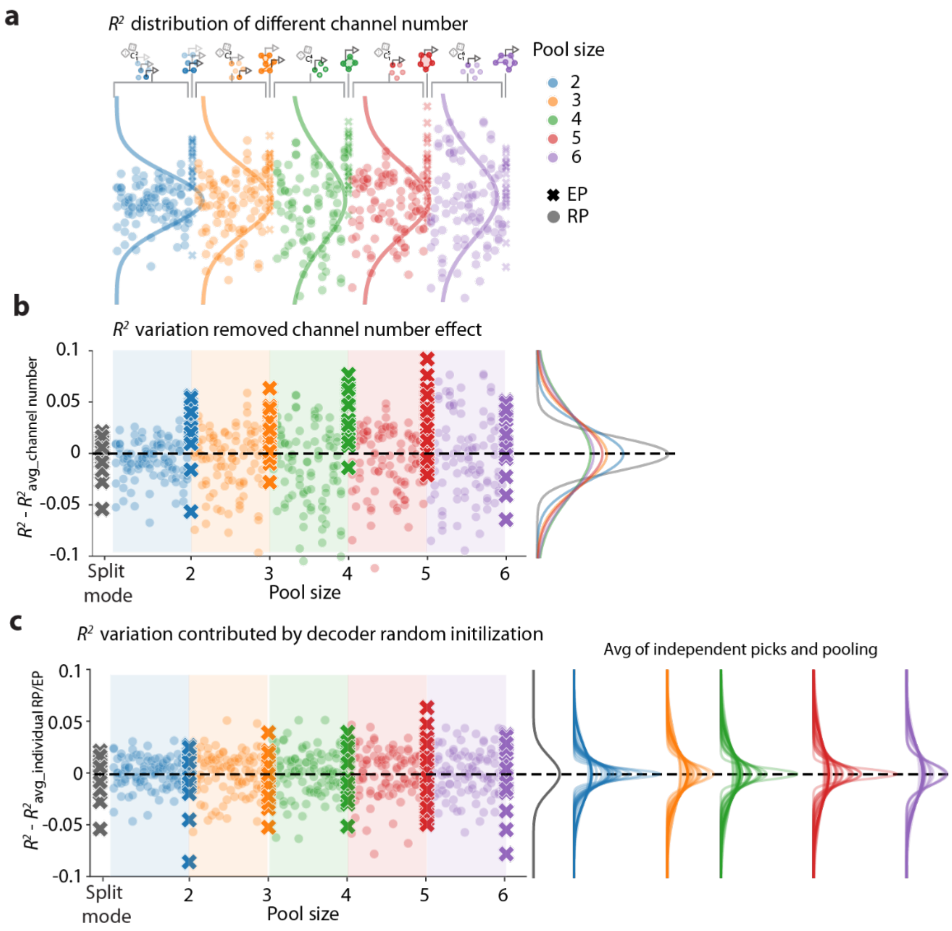


**Fig. S1 | Sources of variability in decoding performance across random-picking and electrode-pooling configurations.**
**a,b,** To isolate variation attributable to channel number, decoding performance was mean-centered within each pool size using the combined electrode-pooling and random-picking results. Specifically, the mean decoding performance was calculated across all electrode-pooling and random-picking data points for each pool size (denoted by color), yielding $R^2_{\text{avg\_channel number}}$, which describes the mean decoding performance under the same channel number. For split mode and electrode pooling, each column shows 25 decoder runs (fivefold cross-validation × five random initializations). For random picking, each column corresponds to one electrode-selection instance evaluated across the same cross-validation folds. Light dots indicate random picking, colored crosses indicate electrode pooling, and grey crosses indicate split mode. Right, pooled distributions for each pool size. Variance was broadly stable across channel number, although split mode showed lower dispersion and pool size 2 was modestly less variable than larger pools. **c,** To isolate variation attributable to cross-validation configuration and decoder initialization, performances were mean-centered within each configuration. Subsequently, the individual decoding performances within each column were adjusted by subtracting the corresponding mean values, yielding the normalized measure $R^2 - R^2_{\text{avg\_individual RP/EP}}$. The distributions at right show comparable dispersion for electrode pooling and random picking within a given pool size, indicating that most of the additional variability in random picking arises from electrode selection rather than decoder fitting.

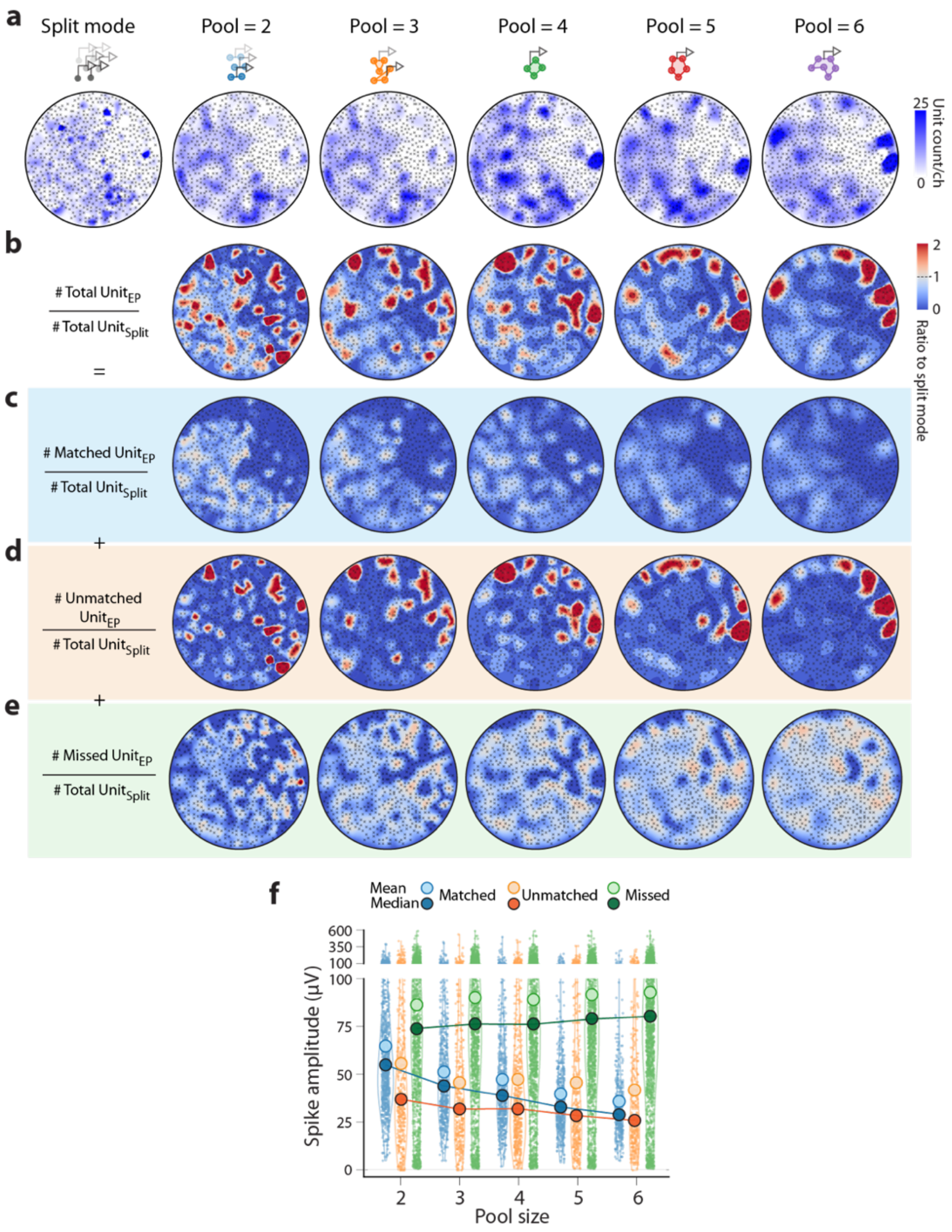


**Fig. S2 | Spatial organization of total, matched, unmatched, and missed unit yields across the electrode array**

**a,** Spatial maps of unit yield on the MwEA (CMOS MEA) in split mode and electrode pooling across pool sizes. **b–e,** Channel-wise ratios of pooled yield relative to split-mode yield for total units, matched units, unmatched units, and missed units, respectively. Pooling increased total yield in the top-right region, but this advantage weakened elsewhere as pool size increased. Matched-unit yields remained comparatively uniform, whereas spatial changes in total yield were dominated by unmatched units. Missed-unit yields were broadly similar to split mode except for channels with zero split-mode yield. **f,** Distributions of negative-peak amplitudes for matched, unmatched, and missed units across channels. Large light and dark circles indicate mean and median, respectively. Median amplitudes of matched and unmatched units decrease with increasing pool size because pooled voltages are averaged across electrodes.

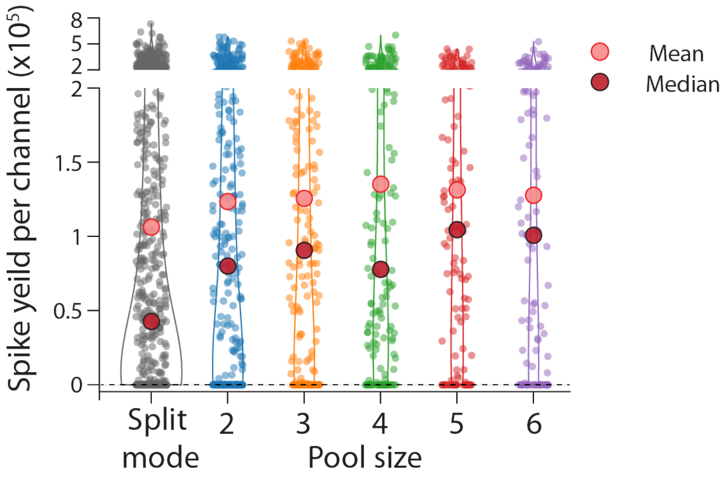


**Fig. S3 | Spike yield per channel increases with pool size**
Violin plots show the number of spikes assigned to each recording channel for pooled configurations. Large light and dark circles indicate mean and median, respectively. Mean spike yield increases with pool size, consistent with aggregation of activity from multiple electrodes onto each transmitted channel.

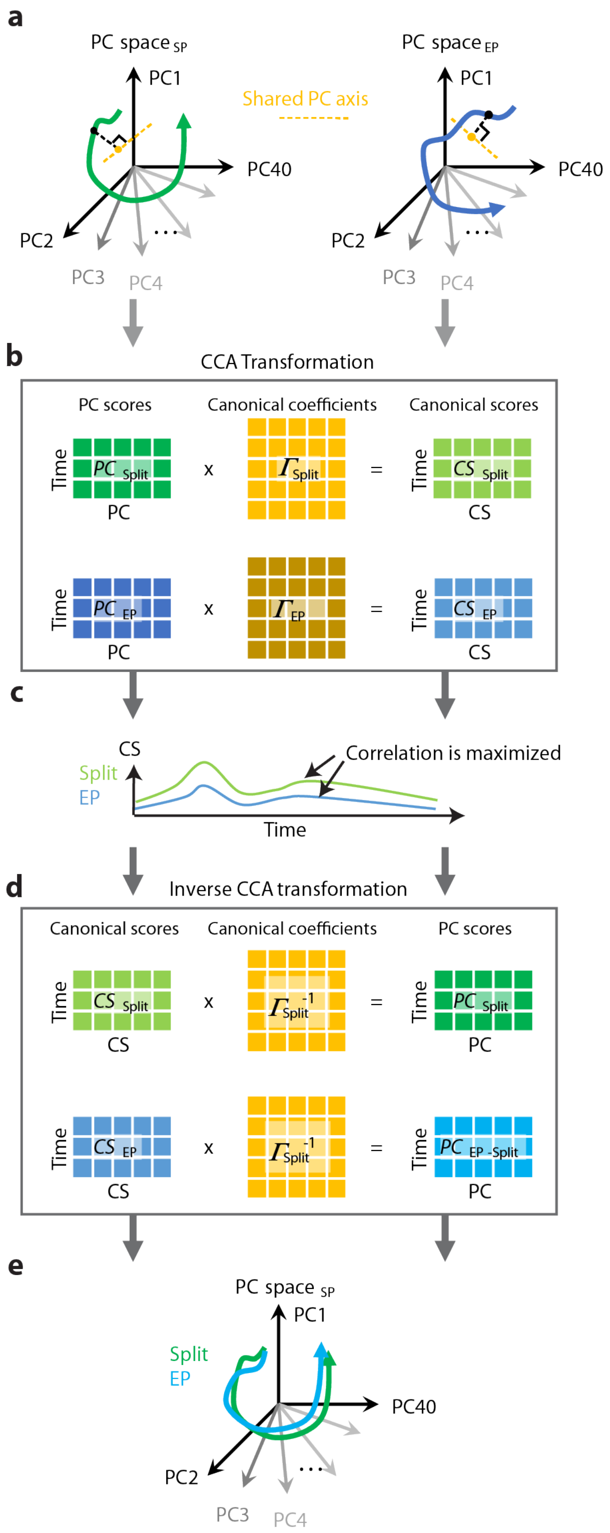


**Fig. S4 | Computational procedure for aligning latent dynamics from pooled and split recordings using canonical correlation analysis**

**a,** Latent dynamics estimated by LFADS were projected into principal component (PC) space. Separate principal component analyses were performed on the split-mode and electrode-pooling latent dynamics, yielding PC score matrices $\boldsymbol{PC}_{\mathrm{Split}}, \boldsymbol{PC}_{\mathrm{EP}} \in \mathbb{R}^{T\times D}$, where $T$ denotes the number of time bins and $D$

denote the number of PCs. **b-c,** Canonical correlation analysis (CCA) was then applied to align the two sets of PC scores within a shared space. Reduced QR decompositions on $\boldsymbol{PC}_{\text{Split}}$ and $\boldsymbol{PC}_{\text{EP}}$ yielded

$$\boldsymbol{PC}_{\text{Split}} = \boldsymbol{Q}_{\text{Split}}\boldsymbol{R}_{\text{Split}}, \boldsymbol{PC}_{\text{EP}} = \boldsymbol{Q}_{\text{EP}}\boldsymbol{R}_{\text{EP}},$$

where $\boldsymbol{Q}_{\text{Split}}, \boldsymbol{Q}_{\text{EP}} \in \mathbb{R}^{T\times D}$ have orthonormal columns ($\boldsymbol{Q}_{\text{Split}}^{T}\boldsymbol{Q}_{\text{Split}} = \boldsymbol{I}, \boldsymbol{Q}_{\text{EP}}^{T}\boldsymbol{Q}_{\text{EP}} = \boldsymbol{I}$) and $\boldsymbol{R}_{\text{Split}}, \boldsymbol{R}_{\text{EP}} \in \mathbb{R}^{D\times D}$ are upper-triangular matrices. Singular value decomposition of $\boldsymbol{Q}_{\text{Split}}^{T}\boldsymbol{Q}_{\text{EP}}$ yielded

$$\boldsymbol{Q}_{\text{Split}}^{T}\boldsymbol{Q}_{\text{EP}} = \boldsymbol{USV}^{T},$$

from which the canonical coefficient matrices for the split-mode and electrode-pooling representations were defined as

$$\boldsymbol{\Gamma}_{\text{Split}} = \boldsymbol{R}_{\text{Split}}^{-1}\boldsymbol{U}, \boldsymbol{\Gamma}_{\text{EP}} = \boldsymbol{R}_{\text{EP}}^{-1}\boldsymbol{V}.$$

Canonical scores were obtained by projecting $\boldsymbol{PC}_{\text{Split}}$ and $\boldsymbol{PC}_{\text{EP}}$ onto manifold directions that maximize pairwise correlations between split-mode and electrode-pooling representations (**c**):

$$\boldsymbol{CS}_{\text{Split}} = \boldsymbol{PC}_{\text{Split}}\boldsymbol{\Gamma}_{\text{Split}}, \boldsymbol{CS}_{\text{EP}} = \boldsymbol{PC}_{\text{EP}}\boldsymbol{\Gamma}_{\text{EP}}.$$

**d-e,** To enable direct comparison of latent dynamics within a shared space, we transformed the canonical scores $\boldsymbol{CS}_{\text{Split}}$ and $\boldsymbol{CS}_{\text{EP}}$ back into the split-mode PC space. Specifically, the mappings were defined as

$$\boldsymbol{PC}_{\text{Split}} = \boldsymbol{CS}_{\text{Split}}\boldsymbol{\Gamma}_{\text{Split}}^{-1}, \boldsymbol{PC}_{\text{EP-Split}} = \boldsymbol{CS}_{\text{EP}}\boldsymbol{\Gamma}_{\text{Split}}^{-1},$$

where $\boldsymbol{PC}_{\text{EP-Split}}$ denotes the electrode-pooling PC scores realigned to the split-mode PC space (**e**). This transformation ensured that both modes were expressed in a shared space, thereby allowing direct comparison of $\boldsymbol{PC}_{\text{Split}}$ and $\boldsymbol{PC}_{\text{EP-Split}}$ for analyzing their latent dynamics.
